# Global landscape of mouse and human cytokine transcriptional regulation

**DOI:** 10.1101/316612

**Authors:** Sebastian Carrasco Pro, Alvaro Dafonte Imedio, Clarissa Stephanie Santoso, Kok Ann Gan, Jared Allan Sewell, Melissa Martinez, Rebecca Sereda, Shivani Mehta, Juan Ignacio Fuxman Bass

## Abstract

Cytokines play a central role in immune development, pathogen responses, and diseases. Cytokines are highly regulated at the transcriptional level by combinations of transcription factors (TFs) that recruit cofactors and the transcriptional machinery. Here, we review three decades of studies to generate a comprehensive database reporting 843 and 647 interactions between TFs and cytokines genes, in human and mouse respectively (http://cytreg.bu.edu). We provide a historic perspective on cytokine regulation discussing research trends and biases. More importantly, by integrating this comprehensive database with other functional datasets, we determine general principles governing the transcriptional regulation of cytokine genes. In particular, we show a correlation between TF connectivity and immune phenotype and disease, we discuss the balance between tissue-specific and pathogen-activated TFs regulating each cytokine gene, and cooperativity and plasticity in cytokine regulation. Finally, we illustrate the use of our database as a blueprint to study TF-cytokine regulatory axes in autoimmune diseases.

## Introduction

Cytokines comprise an array of ~130 polypeptides that are critical in the development of the immune system, and in the regulation of immune and autoimmune responses^1^. Indeed, cytokine dysregulation is associated with myriad diseases including autoimmune disorders, susceptibility to infections, and cancer^1-5^. The expression of cytokines is primarily regulated at the transcriptional level through a combination of tissue-specific (TS) and pathogen- or stress-activated (PSA) transcription factors (TFs)^6,7^. Although cytokine transcriptional regulation has been studied for more than three decades, including hallmark models of transcriptional regulation such as the IFNB1 enhanceosome, we currently lack a comprehensive view of the gene regulatory network (GRN) involved in controlling cytokine gene expression^8^. This limits our understanding of the general principles governing cytokine transcriptional regulation, especially in terms of the relationship between TF connectivity and immune phenotype/disease, the balance between TS and PSA TFs regulating each cytokine gene, and cooperativity and plasticity in cytokine regulation. Here, we review three decades of research to generate a comprehensive and searchable database, CytReg (http://cytreg.bu.edu), comprising 843 human and 647 mouse interactions between TFs and cytokines genes. By analyzing the cytokine GRN and integrating it with phenotypic and functional datasets, we provide novel insights into the general principles governing cytokine regulation and provide a blueprint for further studies.

## Results and Discussion

### Generation of CytReg

To obtain a comprehensive cytokine GRN, we systematically mined ~26 million articles in Medline for studies mentioning at least one of 133 cytokines, one of 1,431 TFs, and an experimental assay (**Fig. 1a**). The resulting 6,878 articles, and 815 additional articles referenced in TRRUST^9^ and InnateDB^10^, were then manually curated to determine whether experimental evidence for the physical and regulatory protein-DNA interactions (PDIs) was provided. This resulted in a list of 1,552 PDIs (843 in human, 647 in mouse, and 62 in other species), for which we annotated the assay used and the regulatory activity identified (**Fig. 1a** and **Supplementary Table 1**). To visualize this GRN we developed a database, CytReg (https://cytreg.bu.edu), where users can browse PDIs by species, TF, cytokine, assay type, and TF expression patterns (**Supplementary Fig. 1**). Links are provided to Uniprot entries for TFs and cytokines, and to PubMed articles reporting the PDIs.

**Figure 1.**
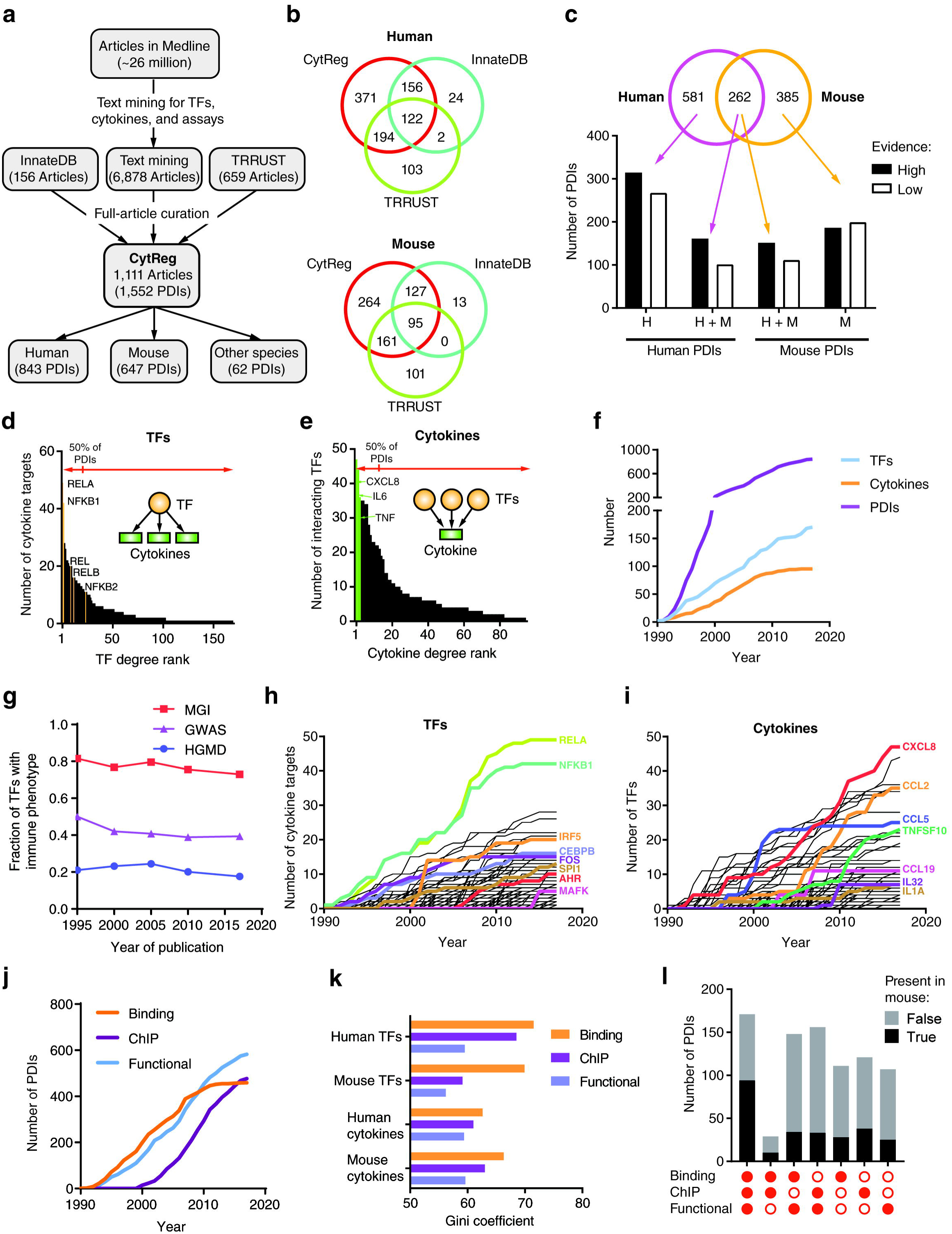
Generation of CytReg and historic perspective of the human cytokine GRN. **(a)** Pipeline used for the text mining and article curation to determine literature-based PDIs between TFs and cytokine genes. **(b)** Overlap of PDIs in CytReg and those annotated in InnateDB and TRRUST. **(c)** Overlap between mouse and human cytokine GRNs (Venn diagram), and fraction of PDIs with high evidence of direct regulatory activity (by a functional assay and an *in vitro* or *in vivo* binding assay) or low evidence (by one type of assay). **(d)** Number of cytokine targets per TF in the human cytokine GRN. **(e)** Number of interacting TFs per cytokine in the human cytokine GRN. **(f)** Number of annotated PDIs, TFs, and cytokines in the human cytokine GRN over time. **(g)** Fraction of TFs in the human cytokine GRN with annotated immune phenotypes when knocked out in mice (MGI) or associated to immune disorders in genome-wide association studies (GWAS) and in the Human Gene Mutation Database (HGMD). **(h-i)** Number of PDIs per TF **(h)** or per cytokine **(i)** in the human cytokine GRN over time. **(j)** Number of PDIs in the human cytokine GRN per assay type over time. **(k)** Gini coefficient to measure the distribution inequality between the number of PDIs per TF or per cytokine. **(l)** Number of PDIs in the human cytokine GRN per assay type and the number of PDIs annotated in the mouse GRN. Filled circles – PDIs involving the assay.

CytReg contains an additional 371 human and 264 mouse PDIs compared to TRRUST and InnateDB (**Fig. 1b**). We also removed 243 PDIs annotated in TRRUST and InnateDB if: a) the article did not provide direct experimental evidence for the PDI, b) the TF interacted with the regulatory region of a cytokine receptor rather than that of a cytokine, or c) the cytokine regulated the activation pathway of a TF rather than the TF regulating a cytokine.

Although multiple PDIs are shared between human and mouse, 69% of human and 60% of mouse PDIs are species-specific (**Fig. 1c**). This low overlap is not likely related to a lack of confidence in the interactions because a similar proportion of interactions found in one or both species were classified as high confidence based on evidence from functional (e.g., reporters assays and TF knockdowns experiments) and binding assays (e.g., chromatin immunoprecipitation -ChIP- and electrophoretic mobility shift assays -EMSAs) (**Fig. 1c** and **Supplementary Table 2**). More likely, this low overlap is related to literature bias and incompleteness of the GRN, or to different modes of regulation between mouse and human.

As observed in other GRNs, a few TFs and cytokines are responsible for most PDIs (**Fig. 1d,e** and **Supplementary Fig. 2**)^11,12^. For example, 12% of the TFs are responsible for more than 50% of the PDIs, including different subunits of NF-κB that when combined represent 16% of the PDIs in the human cytokine GRN (**Fig. 1d**). Similarly, 8% of the cytokines, including the highly studied CXCL8, IL6, and TNF, are involved in more than 50% of the PDIs (**Fig. 1e**). These lopsided distributions in the number of PDIs can be explained by a more central role of some TFs and cytokines in the GRN, but also by research biases as discussed below.

### A historic perspective of the cytokine GRN

By retrieving the publication year of the articles in CytReg, we observed that the size of the cytokine GRN and the number of TFs involved have increased at a constant rate (**Fig. 1f** and **Supplementary Fig. 2**). However, the rate at which new cytokines are incorporated into the network has slowed since 2010 as we are reaching the total number of annotated cytokine genes. This is not the case for TFs as only ~10% of annotated TFs have reported PDIs with cytokine regulatory regions. Importantly, the fraction of TFs that have been incorporated into the cytokine GRN that are associated with immune phenotypes or diseases has remained constant suggesting that the GRN continues to grow towards immune-relevant interactions (**Fig. 1g** and **Supplementary Fig. 2**). Therefore, we expect that the size of this GRN will continue to increase with the development of reagents to test TF functionality.

The time course of PDIs reported per TF differs among TFs (**Fig. 1h** and **Supplementary Fig. 2**). In some cases, such as RELA, NFKB1, and FOS, PDIs were identified since the early 1990s and have increased at a constant rate until the last decade when they plateaued (**Fig. 1h**)^13-15^. This suggests that most of the PDIs involving these highly studied TFs have been identified. For other TFs, such as MAFK, PDIs were discovered much later when an immune role for MAFK was uncovered^16^. While many early studied TFs tend to be involved in more PDIs, this is not always the case. For example, although PDIs involving IRF5 were not reported until the early 2000s^17,18^, IRF5 has more reported PDIs than FOS and CEBPB, whose first reported interactions were in the early 1990s (**Fig. 1h**)^14,19^.

Similarly, the rate at which new PDIs in the cytokine GRN have been reported over time also differs between cytokines (**Fig. 1i** and **Supplementary Fig. 2**). For several highly studied human cytokines, such as CXCL8 and CCL2, PDIs have increased over time with no sign of plateauing (**Fig. 1i**). For other cytokines, such as CCL5 and CCL19, a plateau has been reached with only a few or no additional PDIs in more than a decade (**Fig. 1i**). This plateauing effect could either be related to transient interest in the regulation of such cytokines or because most of the interactions have already been discovered. Given the steep increase in the number of PDIs within a short time-frame, we believe that the former explanation is more likely. As observed for TFs, cytokines whose regulation was studied early (i.e., 1990s) tend to have more PDIs. However, this is not always the case, for example, CCL19 whose first interaction was reported in 2003 has more PDIs than IL1A whose first interaction was reported in 1996 (**Fig. 1i**)^20,21^.

The number of PDIs determined by different methods has increased at different rates. Indeed, we observe nearly a decade lag in the number of PDIs determined by ChIP compared to binding assays (e.g., EMSA) and functional assays (**Fig. 1j** and **Supplementary Fig. 2**), likely because ChIP requires high-quality antibodies and was not adopted by most immunology laboratories until the 2000s. In addition, we observe a plateau for PDIs determined by binding assays in the last decade possibly as a shift away from *in vitro* assays (e.g., EMSAs) towards *in vivo* binding assays such as ChIP, reflecting the increased awareness of the importance of chromatin context in gene regulation (**Fig. 1j** and **Supplementary Fig. 2**). Although historically more PDIs were reported by binding assays than by ChIP, these tend to be less diverse (**Fig. 1k**). This can be explained by binding assays usually requiring both antibodies against a TF, and a known binding sequence within the target promoter for EMSAs or pull-down experiments. Of note, most PDIs were found by two or three methods, usually an *in vitro* or *in vivo* binding assay and a functional assay (**Fig. 1l** and **Supplementary Fig. 2**). PDIs found by one or two methods have been identified at similar rates in the other species (**Fig. 1l** and **Supplementary Fig. 2**).

### Association between TF connectivity and immune phenotype

The differential connectivity between TFs observed in **Figure 1d** can be due to research biases (such as available reagents) and/or a more important role of some TFs in cytokine regulation. Evidence was found for both hypotheses, as highly connected TFs tend to be highly studied (see below), and TFs that bind/regulate multiple cytokine genes also tend to be expressed in immune cells and be associated with immune phenotypes and diseases (**Fig. 2** and **Supplementary Fig. 3**). Indeed, TFs that interact with multiple cytokine genes show higher expression levels in immune cells (**Fig. 2a**) and higher expression enrichment in immune tissues (such as the spleen, bone marrow, and lymph nodes) compared to TFs that interact with only a few or no cytokine genes (**Fig. 2b**). More importantly, highly connected TFs are more frequently associated with immune phenotypes in knockout mouse studies, and with immune disorders as reported in the human gene mutation database (HGMD) and in genome-wide association studies (GWAS) (**Fig. 2c**, **Supplementary Fig. 3**, and **Supplementary Table 3**)^22-24^. For example, the highly connected TF IRF5 is associated with multiple autoimmune diseases, including multiple sclerosis and systemic lupus erythematosus, and leads to low type I interferon, TNF and IL6 production in knockout mice^22-24^. Conversely, the low connected TFs HMGA2, NDS2, and HMBOX1, to our knowledge, have not yet been associated with immune phenotypes or diseases. Overall, these observations highlight the association between TF connectivity and disease, consistent with previous findings in a developmental GRN^25^.

**Figure 2.**
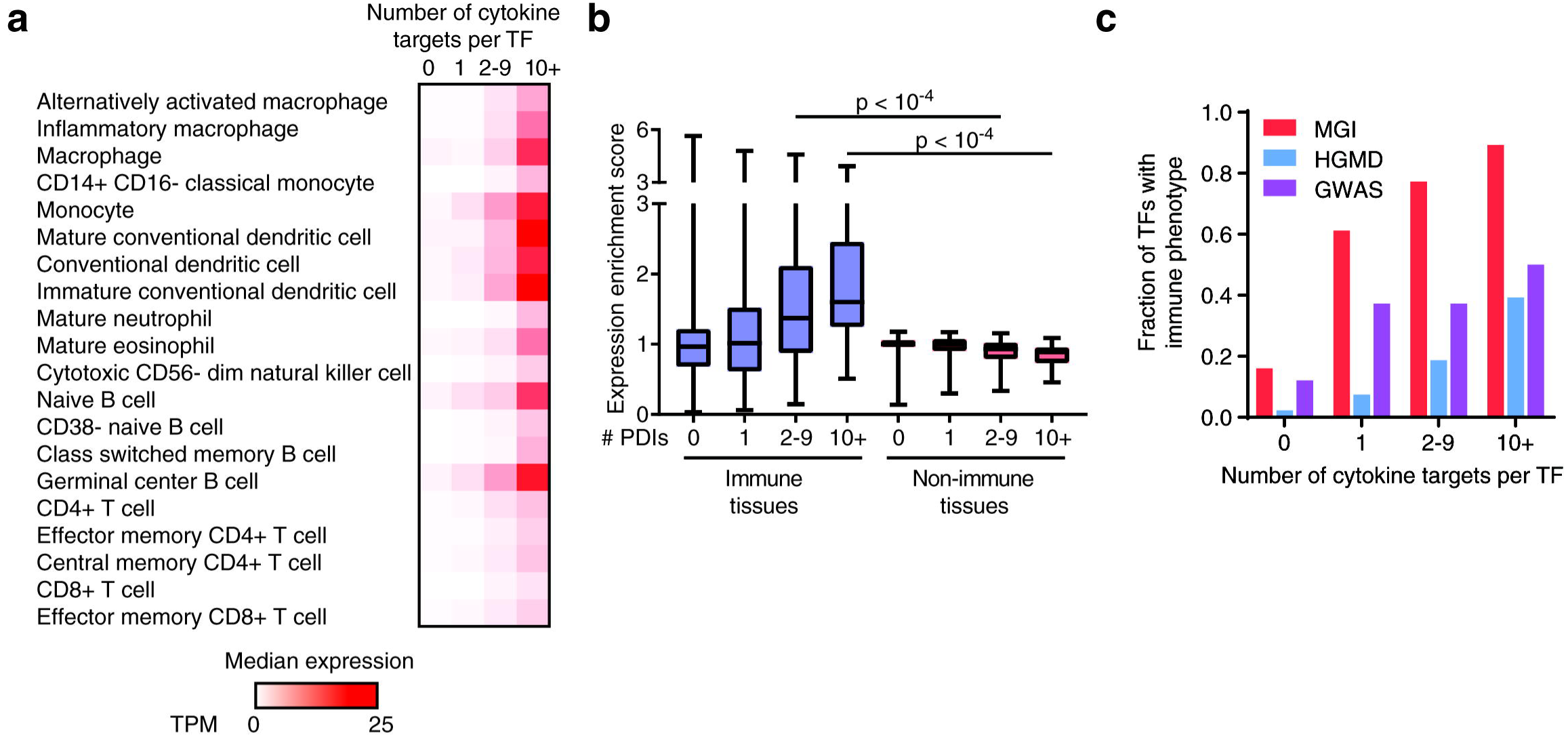
Relationship between TF connectivity and phenotype. **(a)** Median expression as transcripts per million (TPM) across human immune cells obtained from Blueprint Epigenome for TFs displaying different numbers of cytokine targets. **(b)** Expression enrichment in human immune tissues versus non-immune tissues for TFs with varying numbers of cytokine targets. Each box spans from the first to the third quartile, the horizontal lines inside the boxes indicate the median value and the whiskers indicate minimum and maximum values. p-values determined using Wilcoxon matched-pair ranked sign test. **(c)** Fraction of TFs in the human cytokine GRN with annotated immune phenotypes when knocked out in mice (MGI), or associated with immune disorders in the Human Gene Mutation Database (HGMD) or in genome-wide association studies (GWAS) based on the number of cytokine targets.

### Cytokine regulation by different types of TFs

Different cell types express different sets of cytokines in response to pathogen- or cell-mediated cues. For each immune cell type, we determined the TFs enriched in binding/regulating the cytokines expressed in the given cell type (**Supplementary Table 4**). As expected, several master regulator TFs are enriched, including TBX21 (T-bet) in Th1 cells, GATA3 and STAT6 in Th2 cells, RORC in Th17 cells, and SPI1 (PU.1) and CEBPA in monocytes. Additionally, several PSA TFs, such as RELA/NFKB1, are enriched in Th1 cells, monocytes, myeloid dendritic cells, eosinophils, and neutrophils, consistent with these cells producing pro-inflammatory cytokines upon activation; while IRF1/3/5/7 are enriched in B cells and plasmacytoid dendritic cells, producers of type-I interferons in response to viral pathogens.

Highly connected TFs in the cytokine GRN usually belong to the IPT/TIG/p53 (including NF-κB and NF-AT TFs), AP-1, IRF, and STAT families, which are known to play prominent roles in immune cell differentiation and immune responses^26-29^. These TF families are highly enriched in the cytokine GRN compared to the GRN reported in TRRUST^9^, a literature-derived network not constrained to cytokine genes (**Fig. 3a** and **Supplementary Fig. 3**). Furthermore, most PSA TFs are enriched in the cytokine GRN compared to the GRN reported in TRRUST, consistent with many cytokine genes being upregulated in response to pathogens or stress conditions (**Fig. 3b**).

**Figure 3.**
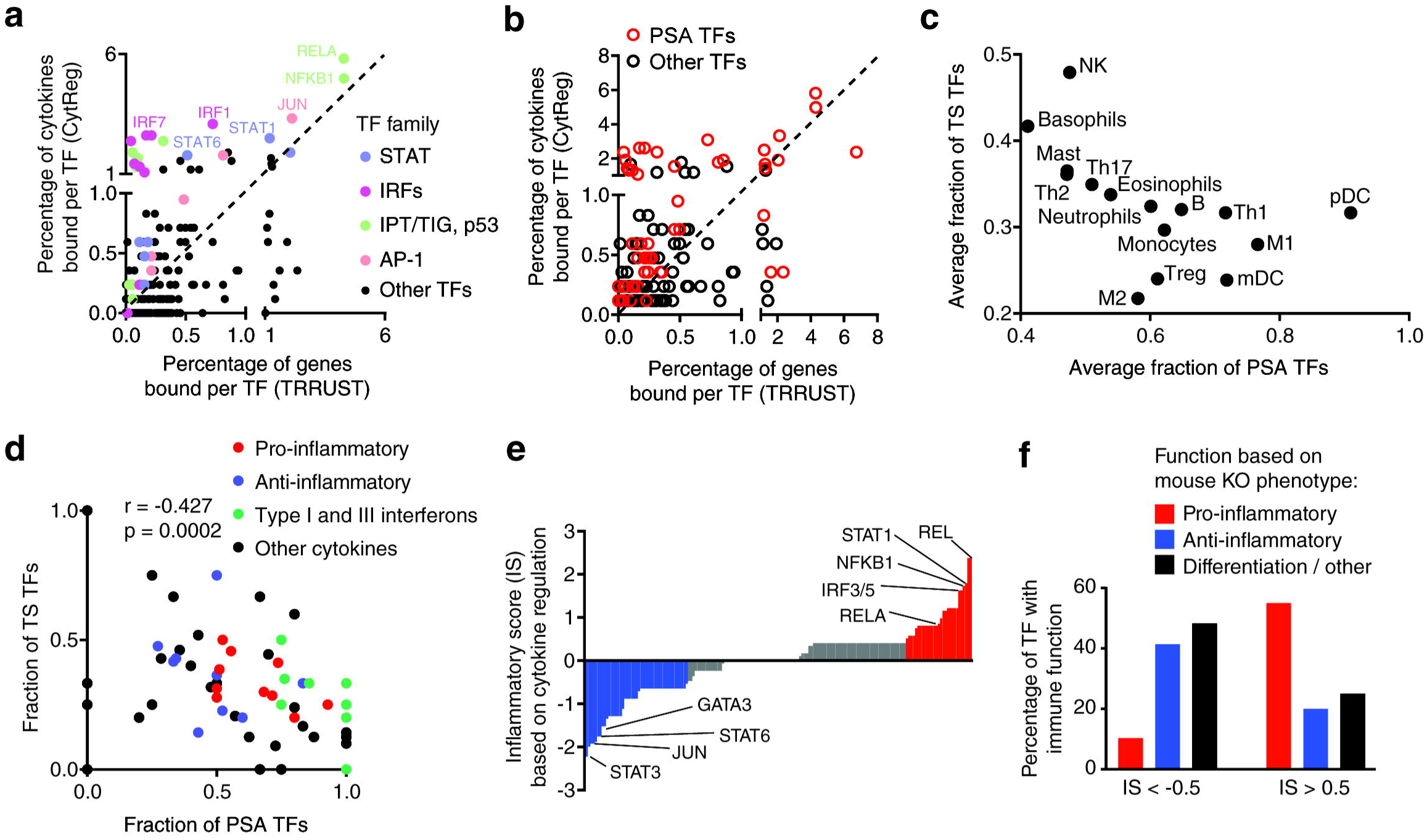
Cytokine regulation by different types of TFs. **(a-b)** Correlation between the percentage of PDIs involving a TF in the human cytokine GRN versus a global human GRN annotated in TRRUST, for different TF families **(a)** or for pathogen- or stress-activated (PSA) TFs **(b). (c)** Average fraction of PSA and tissue-specific (TS) TFs for cytokines expressed in different cell types. **(d)** Fraction of PSA and TS TFs for different classes of cytokines. Correlation determined by Pearson correlation coefficient. **(e)** Inflammatory score (IS) for each TF based on the fraction of PDIs with pro- and anti-inflammatory cytokines. **(f)** Percentage of TFs with pro-inflammatory, anti-inflammatory, and differentiation or other functions based on mouse knockout phenotypes. p = 0.009 by Fisher’s exact test.

Cytokines are expressed in a highly tissue- and condition-specific manner. This is achieved by a specific combination of receptors and signaling pathways present in each cell type, and through the cooperation between PSA and TS TFs^26^. To study the role of PSA and TS TFs in cytokine regulation, for each cytokine we determined the fraction of TFs that respond to pathogen/stress signals (e.g., NF-κB, AP-1 and IRFs) and the fraction of TS TFs based on each TF’s gene expression variability across tissues. Our analysis revealed that cytokines expressed in plasmacytoid dendritic cells, M1 macrophages, Th1 cells, and myeloid dendritic cells are primarily regulated by PSA TFs, whereas cytokines expressed NK cells, basophils, mast cells, Th2 cells, Th17 cells, and eosinophils are also regulated by several TS TFs (**Fig. 3c**). This is consistent with reports of the former cell types expressing multiple canonical pro-inflammatory cytokines and/or interferons, which are induced by pathogen-associated molecular patterns or danger signals from inflammatory microenvironments. Indeed, further analysis revealed that interferons and pro-inflammatory cytokines are regulated by broadly expressed PSA TFs, whereas anti-inflammatory cytokines are regulated by both PSA and TS TFs (**Fig. 3d**).

Different TFs have predominantly pro- or anti-inflammatory functions. Thus, for each TF, we determined an inflammatory score (IS) based on the preference of binding to pro-versus anti-inflammatory cytokine gene targets (**Fig. 3e**). TFs with an IS>0.5 more frequently had a pro-inflammatory function, while TFs with IS<-0.5 more frequently had an anti-inflammatory function based on knockout mouse phenotypes (**Fig. 3f**, p = 0.009 by Fisher’s exact test). Although the dysregulation of other targets is likely involved, these analyses suggest that the cytokine targets of a TF can be important drivers of immune phenotypes.

### GRN integration with TF-cofactor interactions

TFs regulate gene expression by recruiting co-activators and co-repressors that interact with the transcriptional machinery or mediator complex, or that covalently modify histones, TFs, or methylate DNA^30^. Based on literature-derived protein-protein interactions reported in Lit-BM-13^31^, we found that the TFs that bind/regulate cytokine genes interact with numerous cofactors, including multiple co-activators such as EP300, CREBBP, and nuclear co-activators 1-3 and 6 (**Fig. 4a**). This is not surprising given that ~80% of the regulatory PDIs in CytReg are activating and involve potent transcriptional activators such NF-κB and AP-1. Nevertheless, several activating TFs also interact with co-repressors which can inhibit TF function until triggered by signaling pathways^32^.

**Figure 4.**
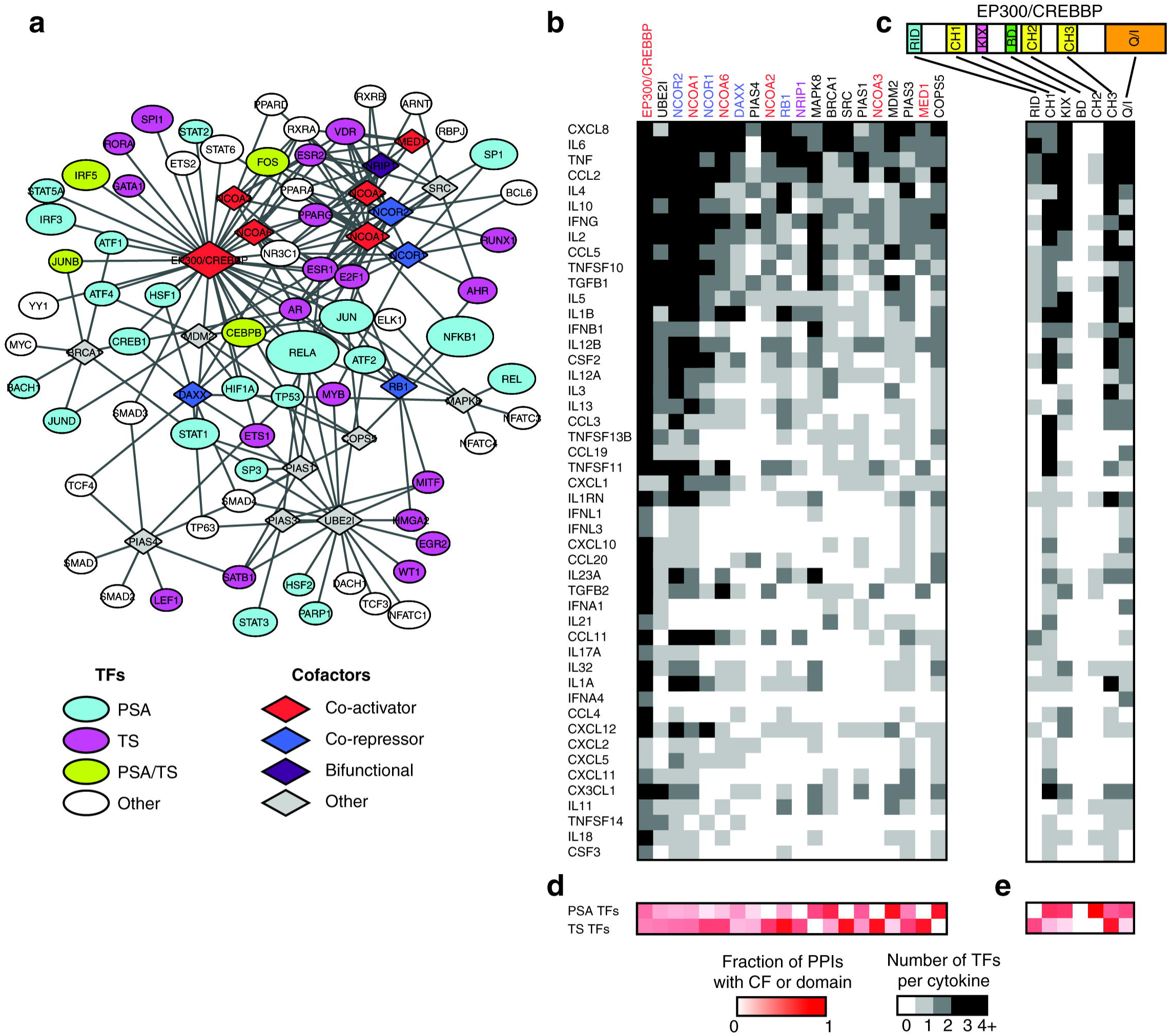
Cooperativity and plasticity in cytokine regulation. **(a)** Protein-protein interaction network from Lit-BM-13 between cofactors and TFs in the human cytokine GRN. Ellipses – TFs, diamonds – cofactors. Node size indicates the number of cytokine targets (for TFs) in the cytokine GRN, and the number of protein-protein interactions with TFs (for cofactors). Only cofactors with five or more protein-protein interactions are shown. **(b,c)** Number of TFs (shades of grey) interacting with each human cytokine gene that interact with the different cofactors **(b)** or the different domains of EP300/CREBBP **(c)**. **(d,e)** Fraction of cofactor **(d)** or EP300/CREBBP domain **(e)** protein-protein interactions (shades of red) involving PSA or TS TFs. Only cytokines and cofactors with five or more interactions are shown. Co-activators are shown in red font, co-repressors in blue font, and bifunctional cofactors in purple font.

In general, each cofactor interacts with multiple TFs that bind/regulate each cytokine gene (**Fig. 4b**)^31^. This may be associated with TF cooperativity to recruit cofactors to regulatory regions as has been reported for the cooperative recruitment of EP300 by RELA, IRFs, JUN, and HMGA1 to the IFNB1 enhanceosome^8^. Alternatively, cofactor binding to multiple TFs may also be associated with regulatory plasticity by which cofactors can be recruited by different sets of TFs to modulate cytokine gene expression in different cell types or conditions. To evaluate these possibilities, we focused on the histone acetyltansferases EP300/CREBBP, which play key roles in immune regulation and differentiation, and whose protein-protein interactions with TFs have been mapped to their different domains^33,34^. We found that, for cytokines for which multiple PDIs have been determined, the set of TFs that bind/regulate that cytokine gene collectively interact with multiple domains of EP300/CREBBP (**Fig. 4c**). This may lead to a cooperative recruitment of EP300/CREBBP to regulatory regions, as has been observed for the IFNB1, TNF, and IL6 genes^8,35,36^. This is also consistent with the observation that, even for cytokines with multiple annotated PDIs, the mutation of a single TF binding site or the inhibition of a single TF can lead to a dramatic effect on gene expression^37,38^. Interestingly, for each cytokine, several TFs can also interact with the same domain of EP300/CREBBP (**Fig. 4c**). Although this may contribute to a cooperative recruitment of EP300/CREBBP, it may also increase regulatory plasticity in different cell types and/or under different stimuli by allowing different TF combinations to induce cytokine expression. For example, TNF induction by LPS, calcium, or viruses all lead to EP300/CREBBP recruitment to the TNF enhanceosome, however, through different sets of TFs^36^.

Some cofactors such as MAPK8, BRCA1, MDM2 and COPS5 preferentially interact with PSA TFs, consistent with their reported function in inflammation and stress responses, and associated immune phenotype in knockout mice (**Fig. 4d**)^23^. Other cofactors such as NCOR1/2, NCOA1/2/3/6, RB1, NRIP1, SRC and MED1 interact primarily with TS TFs such as nuclear hormone receptors^31^. Interestingly, different domains of EP300/CREBBP interact preferentially with PSA or TS TFs: for example, CH1, KIX and Q/I interact mostly with PSA TFs, whereas RID and CH3 interact mostly with TS TFs (**Fig. 4e**). Altogether, this suggests that PSA and TS TFs cooperate in recruiting EP300/CREBBP through different domains to induce cytokine expression under the right stimuli and in the appropriate cell types. In addition, functional redundancy between different PSA TFs may allow for the activation of cytokine expression under different conditions. For example, the PSA TFs HIF1A and NF-κB, both of which interact with the CH1 domain of EP300/CREBBP, can independently induce CXCL8 expression^39^. Overall, these findings are consistent with a model that contains aspects of both the enhanceosome and billboard models of gene regulation, where only certain combinations of TFs present in particular cells or conditions can induce gene expression^40^. Each cytokine, depending on their regulatory flexibility, may be closer to one model or the other.

### The cytokine GRN as a blueprint to study disease

Mutations in multiple TFs have been associated with immune disorders such as autoimmune diseases^22,24^. The role of TFs in autoimmunity is likely related to the dysregulation of immune genes, in particular cytokines, as they play a central role in immune responses and tolerance^4,5^. Indeed, mutations in many cytokine genes have been associated with autoimmunity^22,24^. We considered the cytokines and TFs that have been associated with autoimmune diseases in GWAS and HGMD, and found that many TF-cytokine gene pairs that interact in the cytokine GRN have been associated with the same autoimmune disease (**Fig. 5a**). For example, we found multiple TF-cytokine pairs associated with inflammatory bowel disease, rheumatoid arthritis, atopic dermatitis/psoriasis, and systemic lupus erythematosus (**Fig. 5a** and **Supplementary Table 5**). These TF-cytokine pairs may constitute different regulatory axes by which TFs lead to the disease. For example, AHR activation is protective in inflammatory bowel disease, partly due to increased expression of IL10^41^. Furthermore, TFs associated with autoimmune disorders tend to bind/regulate more cytokines that are themselves associated with autoimmunity (**Fig. 5b**). Overall, the network depicted in **Fig. 5a** may constitute a blueprint to study other regulatory axes in autoimmunity.

**Figure 5.**
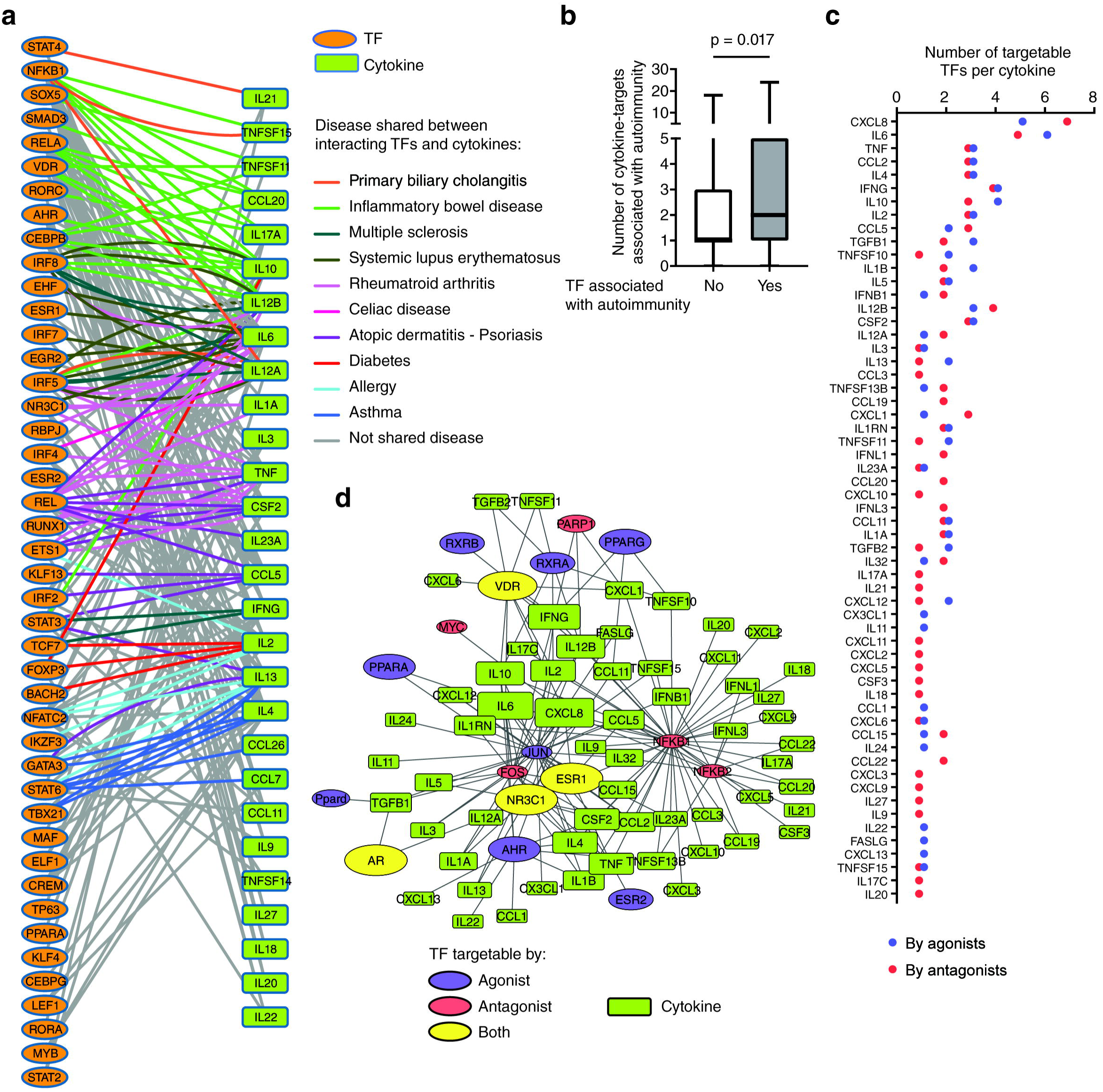
Association of the cytokine GRN with human diseases. **(a)** GRN connecting interacting TFs and human cytokine genes associated with autoimmune or autoinflammatory disorders. Edges connect interacting cytokine-TF pairs. Edge color indicates that the interacting cytokine and TF are associated with the same disease based on HGMD and GWAS. **(b)** Number of cytokine targets associated with autoimmunity for TFs that are (Yes) or are not (No) associated with autoimmunity themselves. Each box spans from the first to the third quartile, the horizontal lines inside the boxes indicate the median value and the whiskers indicate minimum and maximum values. p-value determined by Mann-Whitney U test. **(c)** Number of TFs per cytokine gene that can be targeted by agonists or antagonists. **(d)** GRN connecting cytokines with TFs that can be targeted by approved drugs. Blue, red, and yellow ovals indicate TFs targetable by agonists, antagonists, or both, respectively. Oval size corresponds to the number of approved drugs targeting a TF. Rectangles indicate cytokine genes. Rectangle size is proportional to the number of drugable TFs per cytokine.

Targeting cytokine activity is a widely used therapeutic approach for multiple autoimmune and inflammatory diseases^42,43^. However, currently only ~15% of cytokines can be directly targeted with approved small molecules or specific antibodies, as reported in Drugbank^43^. An alternative strategy is to modulate cytokine production by activating or repressing TF regulatory pathways or by using TF agonists or antagonists^32,43,44^. Although the use of antibodies is a more specific therapeutic approach to inhibit cytokine activity, antibodies cannot be used in many cases because: 1) approved antibodies blocking cytokine activity are only available for nine cytokines, 2) a therapeutic strategy may require the concomitant modulation of multiple functionally related cytokines, or 3) a strategy may require induction of cytokine activity rather than inhibition. In these cases, modulation of cytokine expression by targeting TFs may provide an effective alternative approach.

Many cytokines can potentially be targeted using drugs against their interacting TFs (**Fig. 5c**). Indeed, multiple TF agonists and antagonists have been approved as therapeutics, in particular those that modulate the activity of nuclear hormone receptors (**Fig. 5d**). Targeting these TFs can increase or decrease cytokine expression. For example, IL10 expression can be induced using AHR agonists as a protective mechanism in inflammatory bowel disease, or repressed by an endogenous VDR agonist (calcitriol) during pregnancy to enhance responses to microbial infections^41,45^.

Classic examples of modulating cytokine expression through their interacting TFs include the inhibition of IL2 expression via blocking NF-AT activation to prevent organ transplant rejections, and the inhibition of TNF production in sepsis using proteasome inhibitors to block NF-κB translocation to the nucleus^44,46^. However, the downside of these approaches is that these drugs tend to have multiple side effects since the inhibition of signaling pathways can affect other TFs and biological processes, and each TF can regulate hundreds/thousands of genes in multiple cell types^32,47^. A solution would be to inhibit cytokine expression more specifically by targeting TFs with few interactions in the GRN (less pleiotropic) and by inhibiting TF activity in a more specific manner, for example, by using small molecules that inhibit PDIs or specific protein-protein interactions between TFs and/or cofactors^48,49^. An alternative strategy is to partially inhibit specific combinations of synergistic TFs to reduce the number of potential secondary targets. However, the identity of synergistic TF-pairs regulating each cytokine gene remains to be determined.

### Completeness of the cytokine GRN and future directions

The number of PDIs and TFs in the cytokine GRN have increased at a relatively constant rate over time (**Fig. 1f**), suggesting that the network is still incomplete. There is also a bias towards highly studied TFs and cytokines as we observed a strong correlation between the number of publications in Medline associated with a cytokine or TF and the number of PDIs in the cytokine GRN (**Fig. 6a,b** and **Supplementary Fig. 4**). An argument can be made that highly connected TFs have more pleiotropic functions and thus, are more frequently studied. However, more than 200 TFs absent in the cytokine GRN lead to an immune phenotype when knocked out in mice, many of which are associated with alterations in cytokine expression (**Supplementary Table 3**)^23^. This suggests that many TFs are absent from the cytokine GRN and that many PDIs involving infrequently studied TFs are missing. Indeed, using yeast one-hybrid assays, motif analyses, and luciferase assays we determined that SPIC, a TF absent from the cytokine GRN, interacts with the promoters of CCL23, CCL24, IFNA2, and IL18, showing that the cytokine GRN comprises other TFs (**Fig. 6c,d**). Interestingly, SPIC is involved in the differentiation and function of splenic red pulp macrophages and bone marrow macrophages, producers of the target cytokines^50-54^. Furthermore, even interactions involving frequently studied TFs may be missing from the cytokine GRN, as we also detected known and novel PDIs involving REL (**Fig. 6e,f**). Altogether, this shows that the cytokine GRN is likely to include other TFs and PDIs currently missing from the network.

**Figure 6.**
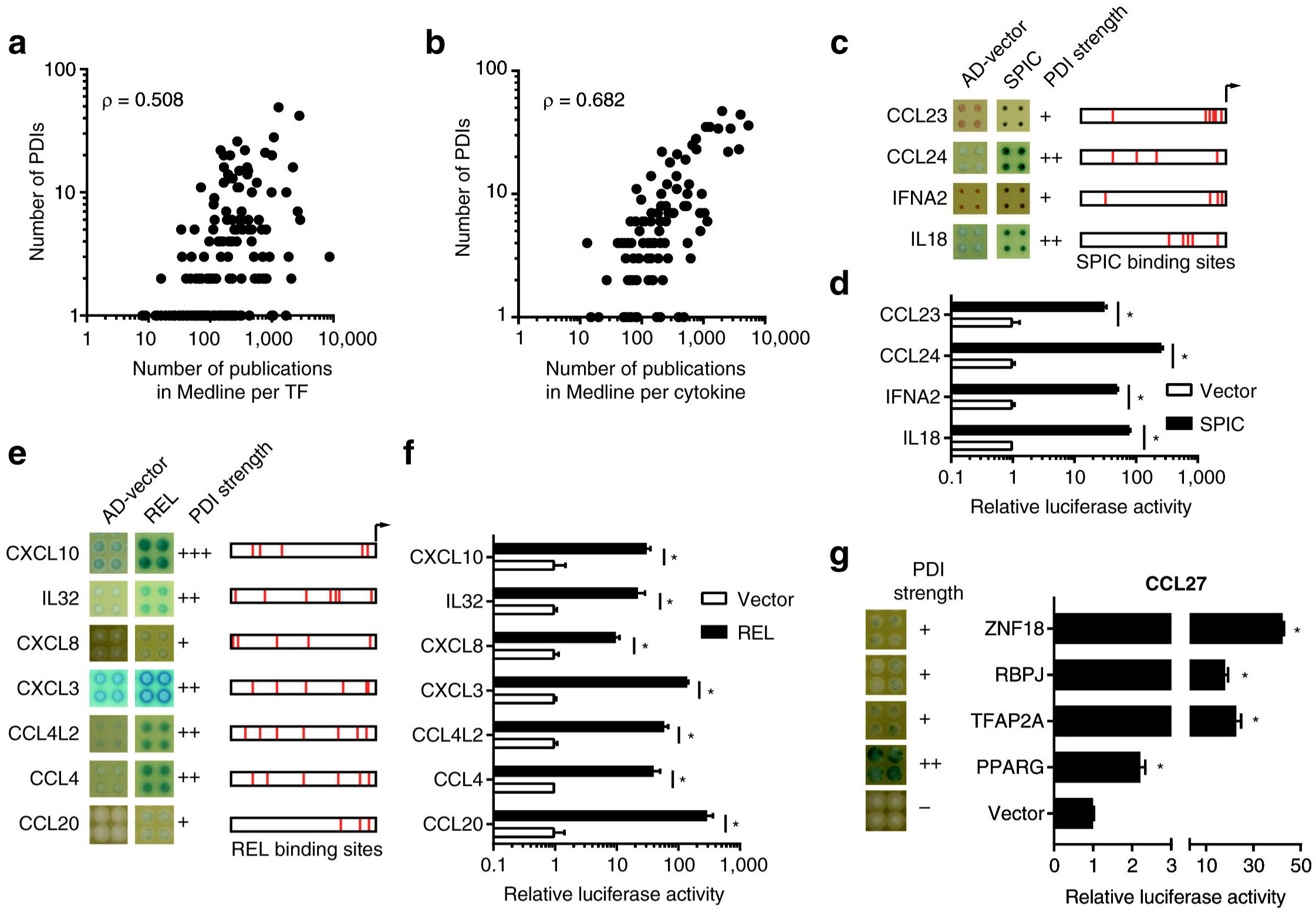
Completeness of the cytokine GRN. **(a, b)** Correlation between the number of PDIs in the human cytokine GRN and the number of publications per TF **(a)** or per cytokine **(b)** reported in Medline. **(c, e)** Enhanced yeast one-hybrid assays testing PDIs between the indicated human cytokine promoters and SPIC **(c)** or REL **(e)**. AD-vector corresponds to empty vector. The qualitative strength of PDIs compared to AD-vector control are indicated as –, +, ++, and +++ corresponding to no, weak, medium, and strong interaction, respectively. REL or SPIC binding sites are indicated in red for each 2 kb promoter region. **(d, f)** HEK293T cells were co-transfected with reporter plasmids containing the cytokine promoter region (2 kb) cloned upstream of the firefly luciferase reporter gene, and expression vectors for the indicated TFs (fused to the activation domain 10xVP16). After 48 h, cells were harvested and luciferase assays were performed. Relative luciferase activity is plotted as fold change compared to cells co-transfected with the vector control (1.0). Experiments were performed in triplicates. Average relative luciferase activity ± SEM is plotted. **(g)** PDIs with the promoter of CCL27 analyzed by eY1H assays (left) and luciferase assays in HEK293T cells (right). *p<0.05 by Student’s t-test.

Similarly, highly studied cytokines are involved in more PDIs (**Fig. 6b**). Although we cannot rule out the possibility that highly studied cytokines have more pleiotropic roles and are regulated by different TFs in different cells and conditions, this alone cannot explain that there are no PDIs reported for 30% of the cytokines. Further, if there is a strong selective pressure to have multiple modes of regulation for certain cytokines, we would expect the mouse and human cytokine orthologs to be regulated by a similar number of TFs, but this is frequently not the case (**Supplementary Fig. 4**). What is more likely is that highly studied cytokines such as TNF and CXCL8 have more PDIs because they have been studied in more cell types and conditions. To test this hypothesis, we performed yeast one-hybrid assays to evaluate the binding of 1,086 human TFs to the promoter of CCL27, a cytokine absent from the GRN (**Fig. 6g**). We detected four interactions involving ZNF18, RBPJ, TFAP2A, and PPARG which were further validated by luciferase assays in HEK293T cells (**Fig. 6g**). Of note, TF ZNF18, which is widely expressed in immune cells, is also absent from the cytokine GRN.

In addition to missing PDIs in the cytokine GRN, individuals may carry genomic variants in noncoding regulatory regions of cytokine genes or in TF coding sequences that lead to different TF-cytokine interactions. Indeed, several single-nucleotide variants (SNVs) have been identified in the promoters of cytokine genes that are associated with diseases that lead to a gain and/or loss of PDIs^25, 55-57^. For example, a SNV in the proximal promoter of CCL5 that is associated with atopic dermatitis leads to a gain of PDI with GATA2^25,55^. Additionally, we recently determined that a SNV in the IL10 promoter, associated with protection against severe malarial anemia, leads to a loss of interaction with the repressor ATF3, which potentially causes increased IL10 expression in patients carrying the protective allele^25,58^. These examples illustrate the complexity of the cytokine GRN. Ultimately, the integration of different high-throughput and unbiased approaches will lead to a more comprehensive picture of cytokine regulation in different cell types, conditions, and individuals.

## Acknowledgements

We thank Brian Gregor for assistance in generating the data mining script, and Drs. Thomas Gilmore and Trevor Siggers for critically reviewing the manuscript. This work was supported by a National Institutes of Health grant to J.F.B. (R00 GM114296 from the NIGMS). J.A.S. was supported by an NIH training grant (5T32HL007501-34 from the NHLBI). M.M. was supported by an NSF-REU (BIO-1659605).

## Authors contributions

S.C.P. performed the data mining in Medline. J.I.F.B., A.D.I., K.A.G., J.A.S., M.M., R.S., and S.M. performed the literature curation. S.C.P. and A.D.I. designed the CytReg web tool. J.I.F.B. and S.C.P performed the data analysis. J.I.F.B., C.S.S., and K.A.G. performed the experiments in Figure 6. J.I.F.B. conceived the project and wrote the manuscript with contributions from S.C.P, C.S.S., and J.A.S. All authors read and approved the manuscript.

---

**Supplementary Figure 1.**
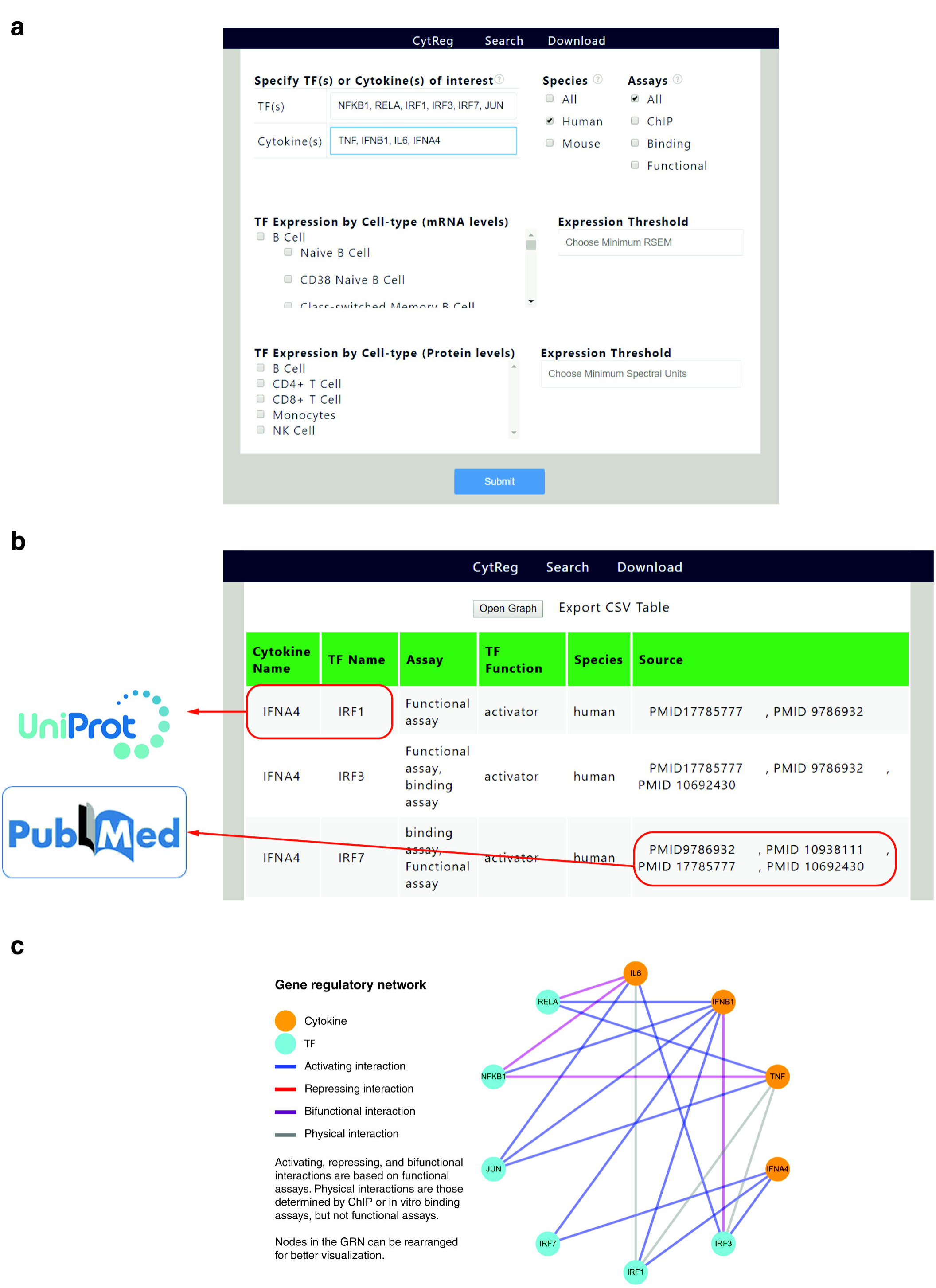

**Supplementary Figure 2.**
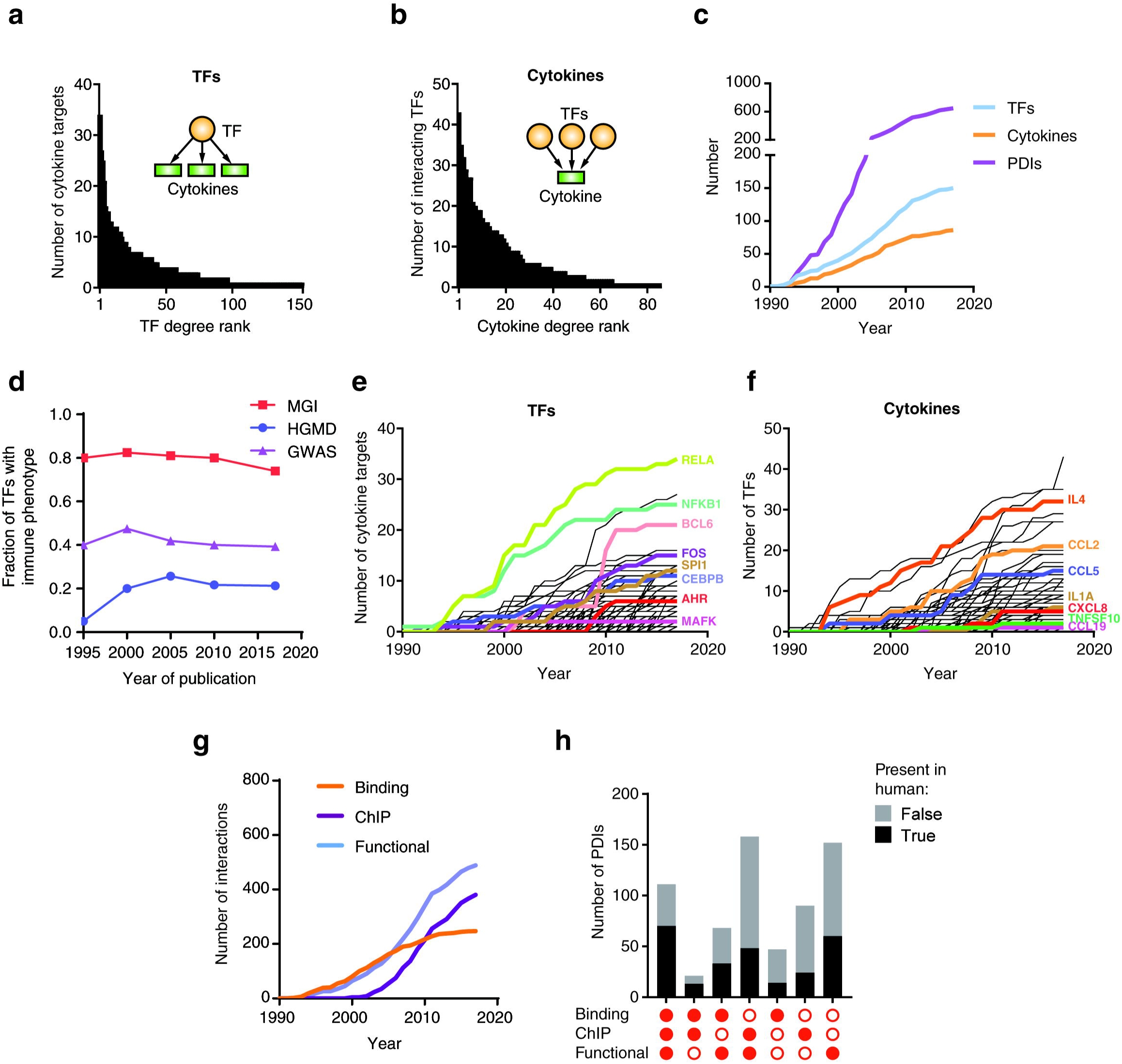

**Supplementary Figure 3.**
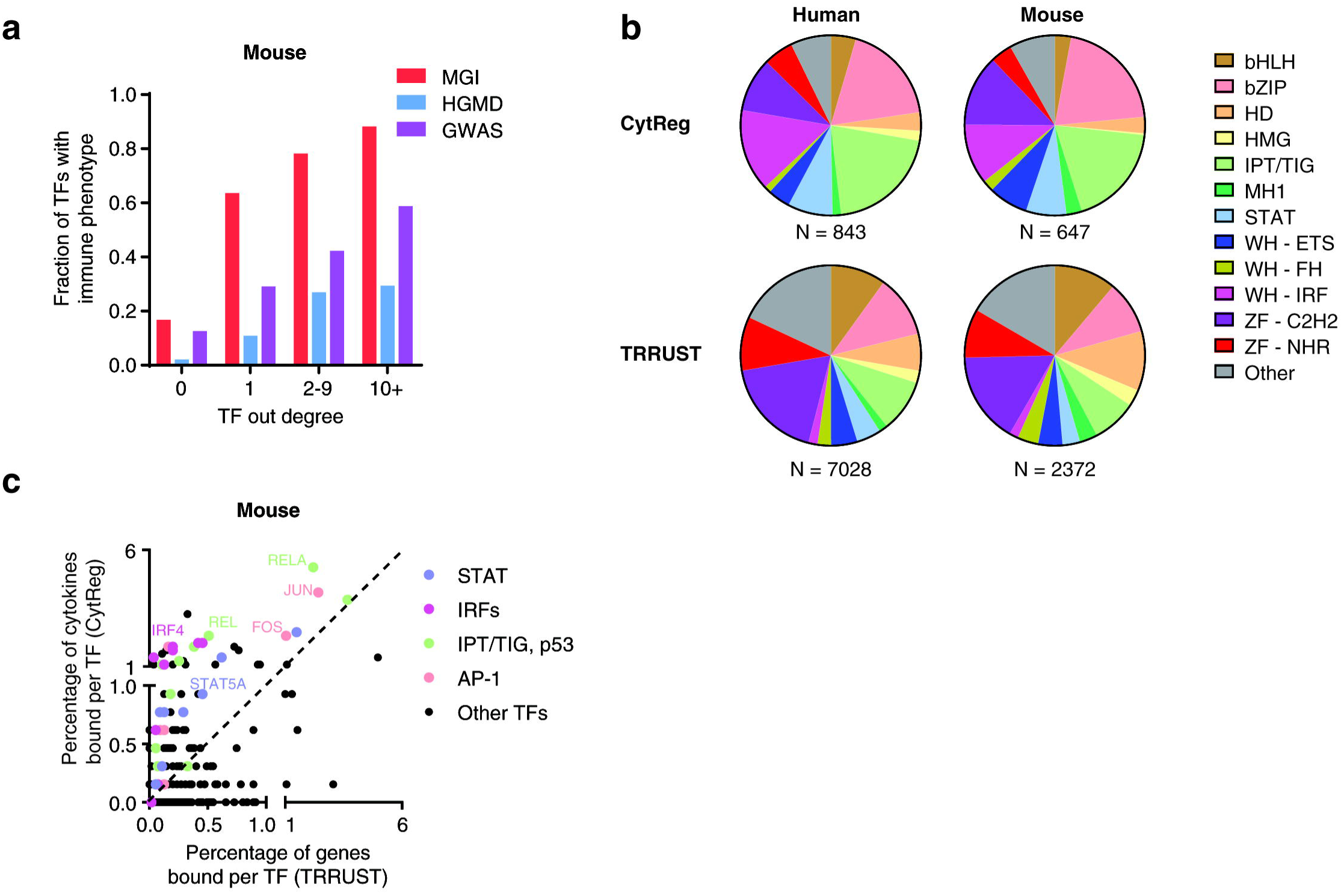

**Supplementary Figure 4.**
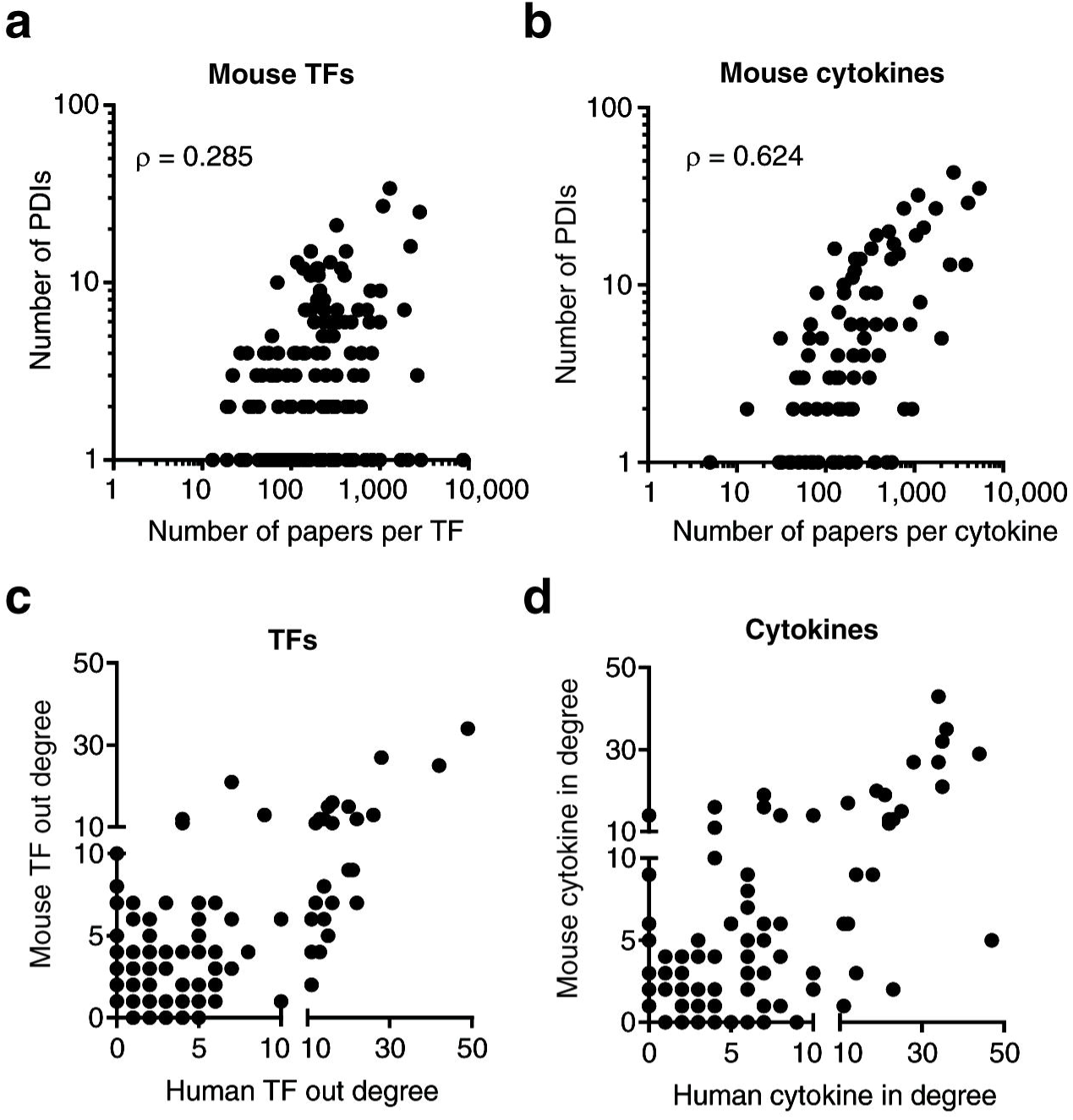

